# In Arabidopsis, low blue light enhances phototropism by releasing cryptochrome 1-mediated inhibition of *PIF4* expression

**DOI:** 10.1101/2020.02.28.969725

**Authors:** Alessandra Boccaccini, Martina Legris, Johanna Krahmer, Laure Allenbach-Petrolati, Anupama Goyal, Carlos Galvan-Ampudia, Teva Vernoux, Elisabeth Karayekov, Jorge Casal, Christian Fankhauser

**Affiliations:** Centre for Integrative Genomics, Faculty of Biology and Medicine, Génopode Building, University of Lausanne, CH-1015 Lausanne, Switzerland; Laboratoire de Reproduction et Développement des Plantes, Univ Lyon, ENS de Lyon, UCB Lyon 1, CNRS, INRAE, 69364 Lyon, France; IFEVA, Facultad de Agronomia, Universidad de Buenos Aires and CONICET, Av. San Martin 4453, 1417 Buenos Aires, Argentina; Fundacion Instituto Leloir, Instituto de Investigaciones Bioquimicas de Buenos Aires–CONICET, 1405 Buenos Aires, Argentina

## Abstract

Shade-avoiding plants including *Arabidopsis thaliana* display a number of growth responses elicited by shade cues including elongation of stem-like structures and repositioning of leaves. Shade also promotes phototropism of de-etiolated seedlings through repression of phytochrome B (phyB) presumably to enhance capture of unfiltered sunlight. Light cues indicative of shade include a reduction in the blue and red portions of the solar spectrum and a low red to far-red ratio. Here we show that in Arabidopsis seedlings both low blue and a low red to far-red ratio are required to rapidly enhance phototropism. However, prolonged low blue treatments through reduced cryptochrome 1 (cry1) activation are sufficient to promote phototropism. The enhanced phototropic response of *cry1* mutants in the lab and in response to natural canopies depends on *PHYTOCHROME INTERACTING FACTORs* (*PIFs*). In favorable light conditions, cry1 limits the expression of *PIF4* while in low blue light PIF4 expression increases, which contributes to phototropic enhancement. The analysis of a quantitative DII auxin reporter indicates that low blue light leads to enhanced auxin levels in the hypocotyl and, upon phototropic stimulation, a steeper auxin gradient across the hypocotyl. We conclude that phototropic enhancement by canopy shade results from the combined activities of phytochrome B and cry1 that converge on PIF regulation.

**ONE SENTENCE SUMMARY:** The persistent depletion of blue light in natural canopy shade relieves the inhibitory effect of cryptochrome 1 on PIF4, enhancing phototropism in de-etiolated Arabidopsis seedlings.

*Financial support:* This work was supported by the University of Lausanne and a grant from the Swiss National Science foundation (n° 310030B_179558 to C.F.); Human Frontier Science Program organization (HFSP) Grant RPG0054-2013, ANR-12-BSV6-0005 grant (AuxiFlo) to T.V.; The University of Buenos Aires (Grant 20020100100437 to J. J. C.), and *Agencia Nacional de Promoción Científica y Tecnológica of Argentina* (Grant PICT-2018-01695 to J. J. C.). Alessandra Boccaccini and Martina Legris are funded by Marie Curie fellowships H2020-MSCA-IF-2017 grants CRoSh 796283 and Flat-Leaf 796443 respectively.

## INTRODUCTION

In natural environments, light conditions are highly dynamic and heterogeneous and given the importance of light for their survival, plants evolved sophisticated photosensory systems to integrate multiple light cues (Casal, 2000; Paik and Huq, 2019). The presence of dense vegetation is not well tolerated by sun-loving plants, such as *Arabidopsis thaliana*. Plants detect neighbors by sensing the low red (R) to far red (FR) ratio (abbreviated as LRFR), which is a consequence of FR reflection by leaves. If the vegetation becomes denser a canopy filters sunlight, creating an environment with LRFR and reduced blue light, red light and Photosynthetic Active Radiation (PAR) (Fiorucci and Fankhauser, 2017). Enhanced hypocotyl elongation and leaf elevation, reduction of branching and flowering acceleration are some of the mechanisms that have evolved to optimize light capture and increase the fitness in response to vegetational shade (Ballare and Pierik, 2017).

Natural canopies are not uniform and gaps allow unfiltered light to create light gradients (Fiorucci and Fankhauser, 2017). Thus, when canopy shade is combined with a directional blue light gradient, plants reorient stem growth to position their photosynthetic organs towards blue light, in a process called phototropism (Ballare et al., 1992; Fiorucci and Fankhauser, 2017). Phototropism is mainly controlled by the phototropin blue light receptors (phot1 and phot2 in *A. thaliana*), which trigger a number of physiological responses including hypocotyl re-orientation towards directional blue light (Liscum and Briggs, 1995; Sakai et al., 2001). Blue light activation of phots generates an asymmetrical distribution of auxin across the hypocotyl, which leads to asymmetrical cell growth between the shaded and lit sides of the hypocotyl (reviewed in (Fankhauser and Christie, 2015)).

Unlike etiolated seedlings, which show high sensitivity to directional blue light, de-etiolated seedlings growing in full sunlight do not show a strong phototropic response (Goyal et al., 2016; Schumacher et al., 2018). However, LRFR, which is typical of shaded environments, enhances phototropism (Ballare et al., 1992; Goyal et al., 2016). The inactivation of phytochrome B (phyB) by LRFR leads to the accumulation/activation of PHYTOCHROME INTERACTING FACTORS 4, 5 and 7 (PIF4, PIF5, PIF7), which promote expression of *YUCCA* genes (*YUC2, YUC5, YUC8*), encoding enzymes for auxin biosynthesis (Hornitschek et al., 2012; Li et al., 2012; Kohnen et al., 2016). This increase of auxin synthesis in the cotyledons promotes hypocotyl re-orientation in LRFR (Goyal et al., 2016).

Phenotypical experiments of seedlings defective for another class of blue light photoreceptors, called cryptochromes (cry), revealed that they modulate phototropism with a positive role in etiolated seedlings (Whippo and Hangarter, 2003; Ohgishi et al., 2004; Tsuchida-Mayama et al., 2010), and a potentially negative role in de-etiolated seedlings (Goyal et al., 2016). The Arabidopsis genome encodes two crys, cry1 and cry2, which coordinate blue light-mediated gene expression by the inactivation of the COP1/SPA E3 ligase complex (Holtkotte et al., 2017; Lau et al., 2019; Ponnu et al., 2019) or through the interaction with several transcription factors (Liu et al., 2008; Ma et al., 2016; Pedmale et al., 2016; Wang et al., 2018; Xu et al., 2018; He et al., 2019; Mao et al., 2020). Light-induced activation of cry1 and cry2 is controlled by BIC1 (Blue light Inhibitor of Cryptochrome 1) and BIC2 (Wang et al., 2016). Cry1 and cry2 are associated with chromatin where they are proposed to control transcription factor activity through incompletely characterized mechanisms (Ma et al., 2016; Pedmale et al., 2016). When expressed in a heterologous system cry2 has the ability to interact with DNA and promote gene expression in a blue light-induced manner (Yang et al., 2018).

At the physiological level crys control several responses such as promotion of blue-light-induced de-etiolation and photoperiodic flowering (Yang et al., 2017). In conjunction with phyB, crys also control shade avoidance responses (Millenaar et al., 2009; Pierik et al., 2009; Keller et al., 2011). In low blue light (LBL) conditions, which is one of the features of canopy shade, the activity of cry is reduced to trigger hypocotyl and petioles elongation (Millenaar et al., 2009; Pierik et al., 2009; Keller et al., 2011; de Wit et al., 2016). One of the mechanisms used by crys to exert their activity is through interaction with and regulation of PIF4 and PIF5 transcription factor activity (Ma et al., 2016; Pedmale et al., 2016). Given the involvement of cry in canopy shade responses and phototropism (Millenaar et al., 2009; Pierik et al., 2009; Keller et al., 2011; Goyal et al., 2016), we decided to determine how crys modulate hypocotyl growth re-orientation in response to blue light features of canopy shade. In our conditions, we found that cry1 is the main cryptochrome involved in the attenuation of phototropism in sunlight-mimicking conditions and the cry1-mediated inhibition of *PIF4* expression is a component of this regulation. Our results reinforce the relevance of the cry1-PIF4 module in light-mediated processes. It emerges as a key module not only for the regulation of hypocotyl elongation, but also for the re-orientation of hypocotyls to avoid canopy shade.

## RESULTS

### Persistent low blue light promotes phototropism

Multiple features of the light environment are altered by canopy shade which can be mimicked in the lab by combining LBL and LRFR (de Wit et al., 2016). In a previous publication, we showed how LRFR enhances phototropism through inactivation of phyB. However, the phenotype of *cry1* suggested that LBL typical of canopy shade also influences hypocotyl re-orientation (Goyal et al., 2016). To determine how specific features of canopy shade (CS) contribute to enhanced phototropism, we measured hypocotyl curvature in the lab under full light (SUN, high blue light and high RFR), under LBL (seedlings were covered up by yellow filter), under low LRFR (FR from the top was added to the white light) or under the combination of both LBL and LRFR as a complete mimic of CS (Figure 1A) (de Wit et al., 2016). Given that LBL enhances hypocotyl growth more slowly than LRFR (Pedmale et al., 2016) we decided to include treatments with different light qualities 24h prior to testing their phototropic potential (LRFR/LRFR and LBL/LBL, Figure 1A). In all conditions analyzed, the seedlings were exposed to supplementary horizontal blue light (8 μmol m^−2^s^−1^) during phototropic stimulation (Figure 1A). We measured deviation from vertical growth after 6 hours of lateral blue light treatment. The overall bending of WT (Col-0) seedlings in SUN/SUN, SUN/LBL and SUN/LRFR was modest, indicating that neither LBL nor LRFR alone were sufficient to trigger a significant enhancement of hypocotyl curvature (Figure 1B,C). However, in SUN/LBL seedlings showed the tendency to bend more and phototropism was significantly enhanced when LBL was combined with LRFR (SUN/CS) (Figure 1B,C). The LRFR condition described in Goyal et al., 2016, did stimulate phototropism; however, here seedlings were grown in long-days under stronger white light, to more closely mimic a natural environment. Interestingly, LBL starting during the pretreatment period enhanced the phototropic response at equal conditions during the exposure to the BL gradient (compare LBL/LBL vs SUN/LBL, Figure 1). In the presence of the same amount of blue light provided unilaterally, the yellow filter used to create the LBL environment changes the blue light differential between the top and the illuminated side. However, this does not appear to be enough to affect phototropism and allowed us to specifically study the effect of LBL on phototropic responsiveness (Figure 1B, and further experiments below). Specifically, LBL but not LRFR pre-treatment affected the phototropic response analyzed on the second day of the experiment (Figure 1B,C; Supplemental Figure S1A), although both treatments induced hypocotyl elongation (Supplemental Figure S1B). Moreover, a treatment with a neutral filter to reduce PAR intensity the day before phototropic stimulation did not affect phototropism (Supplemental Figure S1C). To better define when the LBL pre-treatment was most effective to promote phototropism the following day, LBL treatment was started or ended at different times of the first day (Supplemental Figure S1D). This experiment showed that to be effective the LBL treatment had to occur during the last 4-7 hours of the day prior to phototropic stimulation (Supplemental Figure S1D). We therefore conclude that specifically a prolonged reduction of blue light in the environment promotes phototropism and this is not merely a consequence of enhanced hypocotyl elongation.

**Figure 1.**
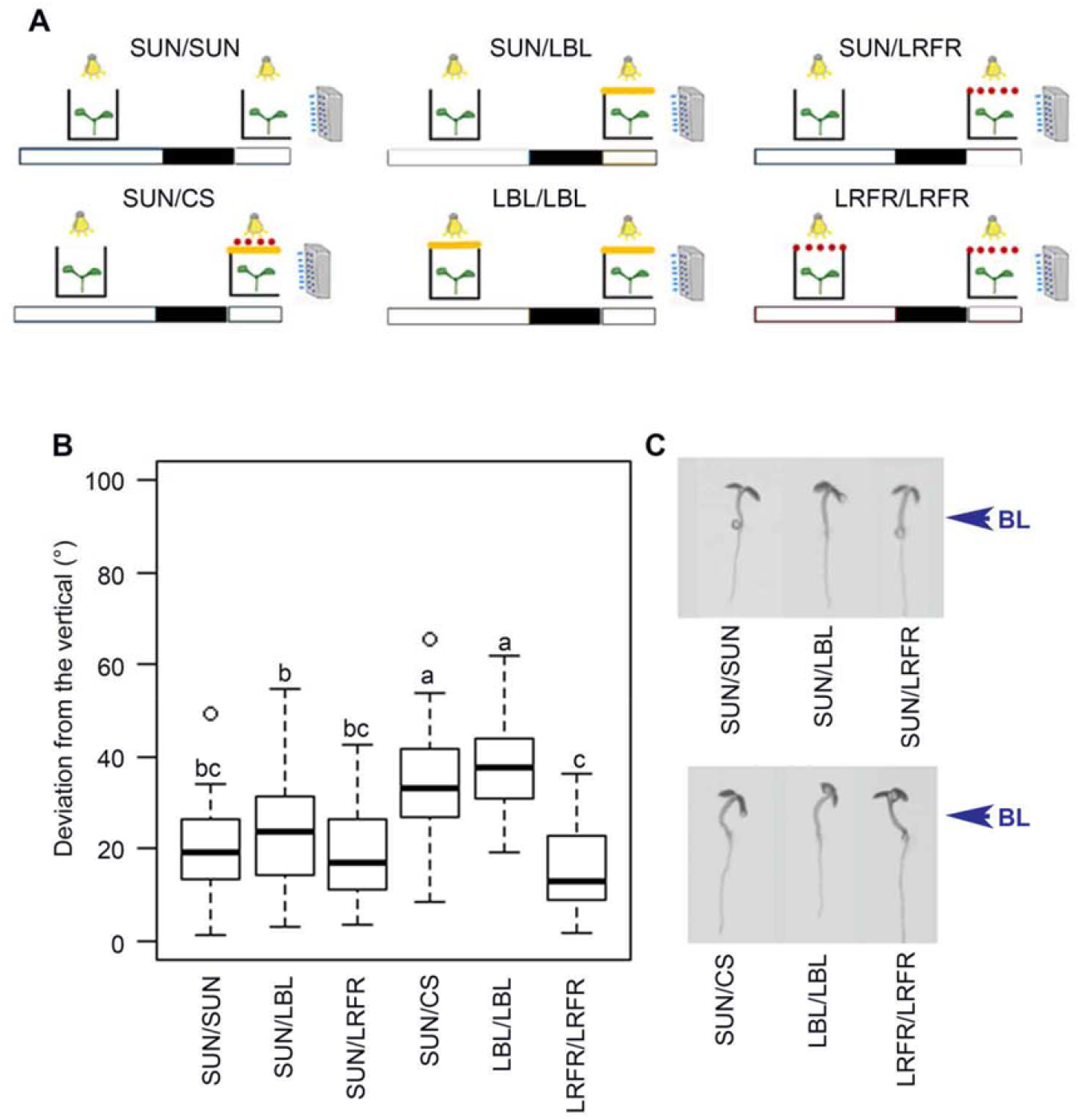
The blue light component of canopy shade is critical for phototropism in green seedlings. A) Experimental scheme, which represents the day preceding the application of lateral blue light and treatment during phototropism (SUN, High Blue light/High Red Far Red; LBL, Low Blue Light; LRFR, low R/FR). Bulbs represent the sources of white light, orange lines represent the filters used to lower BL, red dots represent the sources of FR and blue dots represent the sources used to provide horizontal BL. At ZT0 the light conditions were the same as the previous day, the new light treatment was started within a few minutes after ZT0. For irradiance values refer to Materials and Methods. B) Box plots represent the deviation from the vertical of 4-days old seedlings (n≥25) 6h after lateral blue light application. Letters indicate statistically significant differences at p-value < 0.05 obtained by two way ANOVA followed by the post hoc Tukey’s HSD. C) Representative seedlings of experiment shown in panel B.

### Persistent LBL relieves the inhibitory effect of cry1 on phototropism

Cryptochomes are the photoreceptors sensing blue light reduction in canopy shade (Keller et al., 2011; de Wit et al., 2016; Pedmale et al., 2016) and they also modulate hypocotyl re-orientation in etiolated seedlings (Whippo and Hangarter, 2003; Ohgishi et al., 2004; Tsuchida-Mayama et al., 2010). To define cryptochrome function during shade-enhanced phototropism, we compared hypocotyl growth re-orientation of the wild type and *cry1* mutant in response to different SUN and LBL (pre-)treatment combinations (Figure 2A, B). When phototropism was performed in LBL, 24 hours of LBL pre-treatment strongly accelerated the phototropic response of WT seedlings (Figure 2A). Remarkably, *cry1* seedlings were insensitive to the high levels of blue light present under SUN pre-treatment conditions and responded like the LBL-pretreated WT (Figure 2A, Supplemental Figure S2A). Our experiments showed that a LBL pretreatment enhanced phototropism when it was analyzed either in LBL (Figure 2A) or SUN (Figure 2B) showing that the enhanced response is not due to a change in the blue light gradient. To confirm this, we performed the same experiments in SUN conditions but increased the horizontal blue light intensity to match the gradient in LBL (see Materials and Methods). Both in WT and *cry1* we did not detect significant differences between the SUN responses in high versus low gradient (Supplemental Figure S2B). In addition, the increased gradient in SUN never led to the phenotype observed in LBL/LBL (Supplemental Figure S2B). Lastly, in our light conditions, only *cry1* and *cry1cry2*, but not *cry2*, exhibited a de-repressed phototropic response similar to LBL-pretreated WT seedlings (Figure 2C). Taken together our experiments indicate that cry1 suppresses the phototropic response in SUN conditions and reduced cry1 activation in LBL releases this suppression.

**Figure 2.**
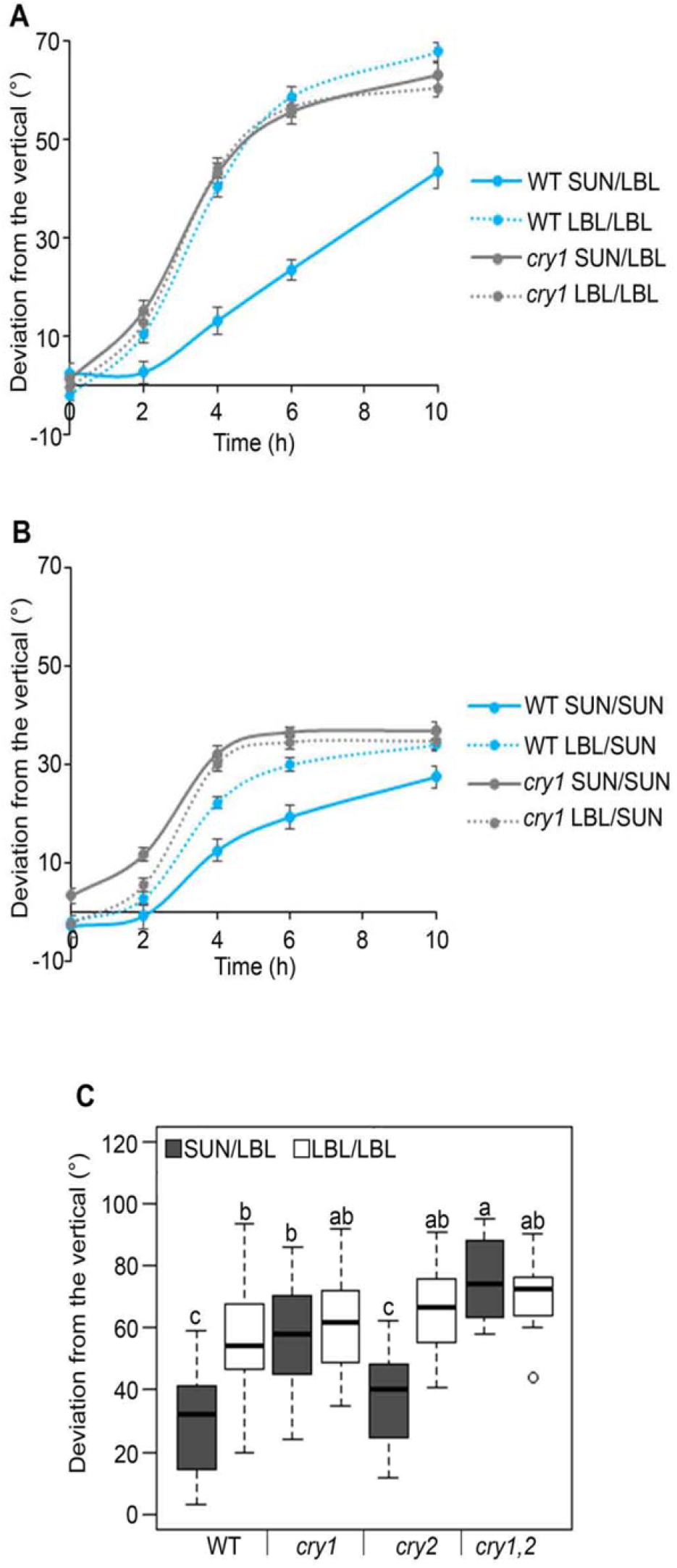
Persistent LBL relieves the inhibitory effect of cry1 on phototropism. Time course analysis of hypocotyl curvature in WT and *cry1* 3-day old seedlings in LBL (A) or in SUN (B) with or without LBL pre-treatment. C) Box plots represent the deviation from the vertical of 3-day old seedlings (n≥25) 6h after lateral blue light application. Letters indicate statistically significant differences at p-value < 0.05 obtained by two way ANOVA followed by the post hoc Tukey’s HSD.

### phot1 is needed for phototropism in LBL

phot1 is the major photoreceptor initiating phototropism towards relatively low blue light intensities both in etiolated and light-grown seedlings (Sakai et al., 2001; Christie et al., 2011; Goyal et al., 2016). Therefore, we assessed whether phot1 was also involved in LBL and cry1-modulated phototropism. We compared the response of the *phot1cry1* with *cry1* and *phot1* single mutants (Figure 3A). *phot1cry1*, as well as *phot1* hypocotyls re-oriented much less than WT in persistent LBL (LBL/LBL), indicating that phot1 was needed for cry1-mediated phototropism enhancement. Moreover, NPH3, which is essential for phototropism in etiolated and green seedlings (Motchoulski and Liscum, 1999; Goyal et al., 2016) was also required for the response in our conditions (Figure 3B). One of the first steps in phot1 signaling is NPH3 de-phosphorylation, which was recently implicated in modulating the phototropic response as reduced NPH3 de-phosphorylation correlates with accelerated phototropism in seedlings treated for a few hours with light prior to phototropic stimulation (Sullivan et al., 2019). We therefore tested whether the LBL treatment that accelerates phototropism led to changes in NPH3 phosphorylation. NPH3 immunoblots did not reveal any differences among the tested light conditions suggesting that the differences in hypocotyl curvature triggered by LBL were not a consequence of altered NPH3 phosphorylation status (Figure 3B). We therefore conclude that LBL-enhanced phototropism requires phot1 and NPH3 but we have no evidence for a role of LBL-regulated NPH3 phosphorylation in this process.

**Figure 3.**
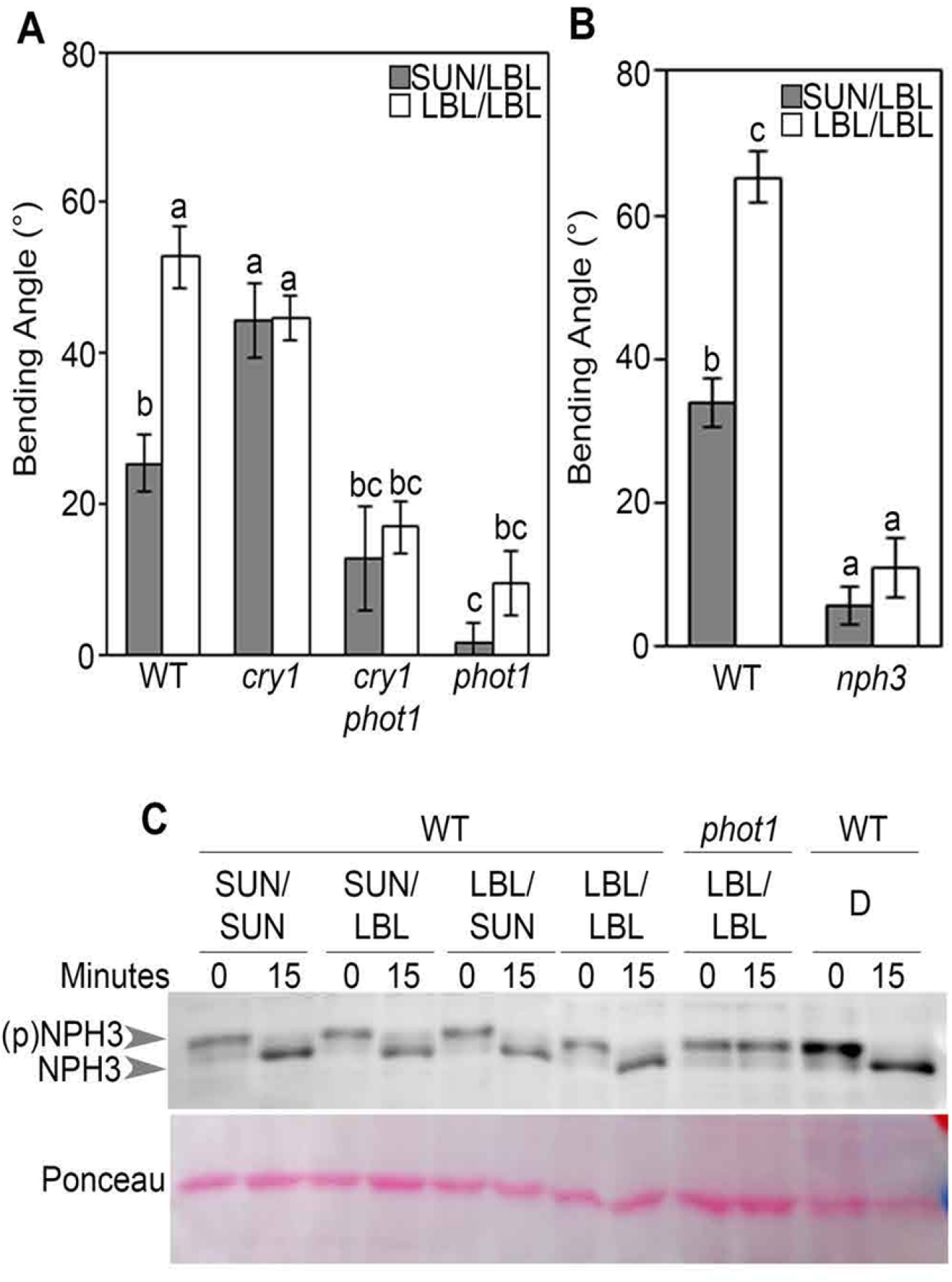
Phototropism in LBL requires phot1. Phototropism in WT, *cry1*, *phot1*, and *cry1phot1* (A) and WT vs *nph3* seedlings (B). All the measurements were done in 3-day old seedlings 6h after lateral blue light application. Letters indicate statistically significant differences at p-value < 0.05obtained by two way ANOVA followed by the post hoc Tukey’s HSD (n≥25). C) Detection of NPH3 phosphorylation state in 3-day old de-etiolated (D) WT and *phot1-5* seedlings before the end of the night (T0) and 15’ (T15’) minutes after dawn in presence of lateral blue light. NPH3 was also detected in WT etiolated seedlings before (T0) and after 15 minutes (T15’) of lateral blue light (8 μmol m^−2^s^−1^). Ponceau staining was used as loading control.

### PIF4 and 5 modulate LBL-dependent phototropism downstream of cry1

Crys act trough PIFs to regulate hypocotyl elongation in response to temperature (Ma et al., 2016) and blue light (Pedmale et al., 2016). To understand if the cry1-PIFs module also operates during shade-controlled phototropism, we analyzed the phototropic bending of different combinations of *cry1* and *pif* mutants in three different light conditions with the same blue light gradient: SUN/LBL, LBL/LBL and SUN-CS. The *pif4pif5pif7* triple mutant had the same phototropic response as the WT in SUN/LBL but showed no phototropism enhancement in response to LBL pretreatment or CS treatment the day of phototropism (Figure 4A). Remarkably, the *pif4pif5pif7* triple mutant was epistatic over *cry1* in all tested conditions (Figure 4A). Interestingly, *pif4pif5* double mutants were unresponsive to the LBL pretreatment, while they responded normally to the CS treatment (Figure 4A). Moreover, *pif4pif5* double mutants selectively suppressed the *cry1* phenotype in SUN/LBL and LBL/LBL but not SUN/CS conditions (Figure 4A). To test the relevance of these findings in natural conditions we analyzed the phototropic response outdoors in response to a real canopy (Figure 4B). Seedlings were grown in the lab for 4 days before being placed on the south side of a grass canopy (Southern hemisphere) (Figure 4B). Both *phyB* and *cry1* mutants re-oriented more than WT seedlings, while *pif4pif5pif7* showed a weaker phototropic response (Figure 4B). Interestingly, in this condition, *pif4pif5pif7*, but not *pif4pif5*, fully suppressed the *cry1* phenotype as observed in the lab in SUN-CS conditions (Figure 4A, B). Moreover, while *pif4pif5pif7* was fully epistatic over *cry1*, this triple mutant did not fully suppress the *phyB* phenotype (Figure 4B). Taken together our laboratory and outdoor experiments indicate that PIF4 and PIF5 are specifically required for the LBL response downstream of cry1. In contrast, the response to real canopy shade (LBL and LRFR), also requires PIF7 (Figure 4) (Goyal et al., 2016).

**Figure 4.**
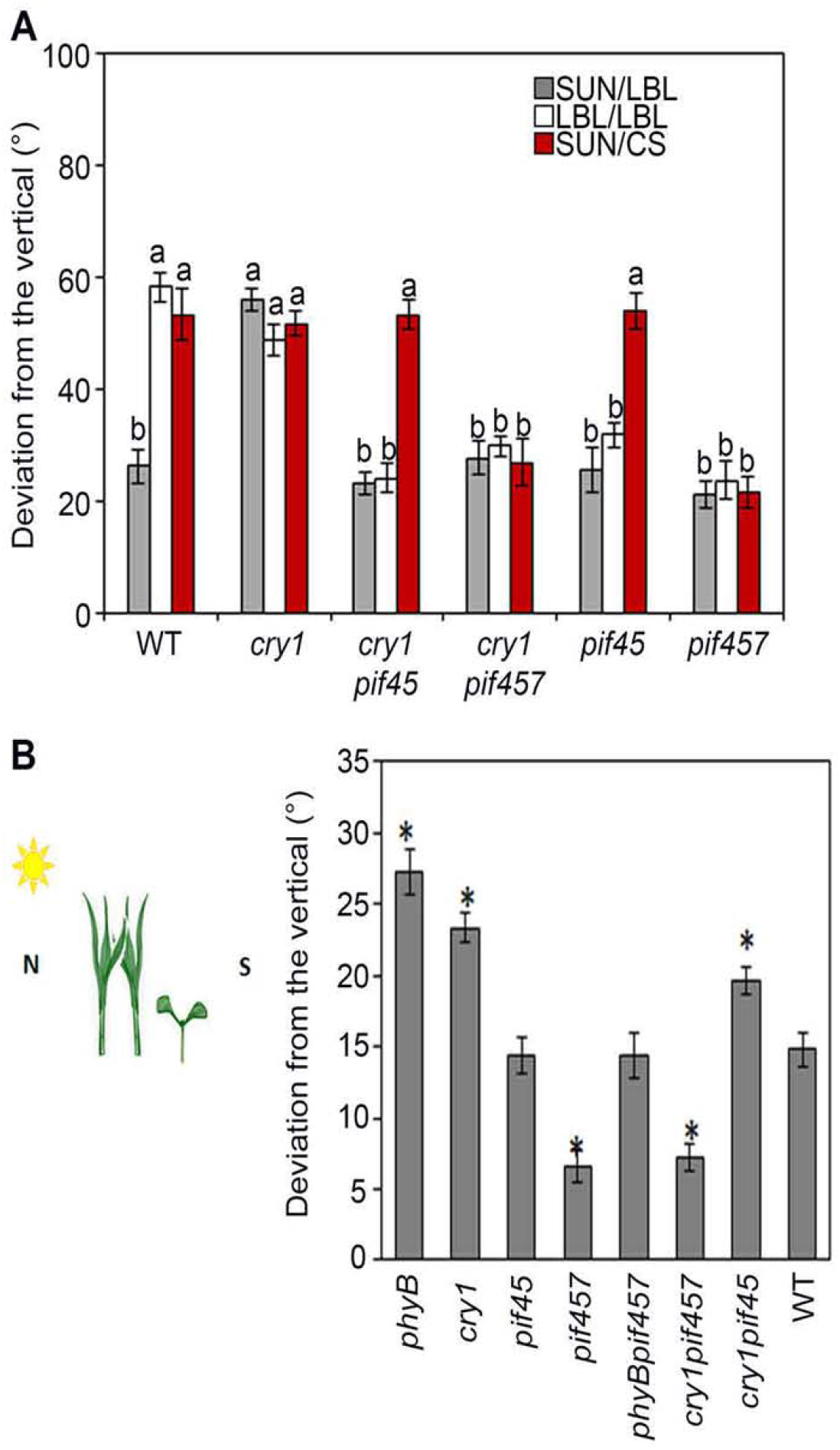
PIF4 and PIF5 act downstream cry1 to control phototropism. A) Bars represent mean values of WT, *cry1, pif4pif5 (pif45), pif4pif5pif7 (pif457), cry1pif4pif5 (cry1pif45), cry1pif4pif5pif7 (cry1pif457)* hypocotyl deviation from the vertical in the lab conditions. Letters indicate statistically significant groups at p-value < 0.05 obtained by two way ANOVA followed by the post hoc Tukey’s HSD. B) Scheme of the outdoor experiment (left). Seedlings were grown for 4 days in lab condition, then were moved on the south side of grass plants for phototropic assay and hypocotyl bending was measured 5hours after phototropic stimulation. Bars represent (right) mean values ±SE of three independent experiments. Asterisks indicate the statistical significance (Student’s t-test p< 0.05) of mutant bending respect to WT.

### LBL enhances PIF4 protein levels towards the end of the day

Given the importance of PIF4 and PIF5 in regulating hypocotyl curvature (Figure 4), we wondered whether the faster phototropic response observed under prolonged LBL (Figure 2B) correlated with a faster accumulation of PIF4 and/or PIF5. We determined PIF4 (Figure 5A, B) and PIF5 (Figure 5A, C) protein levels, using lines expressing the *PIF4/5-HA* transgene under the control of their native promoters (*PIF4p:PIF4-HA* in *pif4* and *PIF5p:PIF5-HA* in *pif5*) during the first 3 hours of the phototropic response in LBL with or without LBL pre-treatment. As reported previously (Bernardo-Garcia et al., 2014; Galvao et al., 2019), levels of both PIF4 and PIF5 increased from ZT0 to ZT3, but we did not observe an effect of the LBL pretreatment on PIF protein levels (Figure 5B, C). Given that LBL-enhanced phototropism is most effective with a LBL pretreatment we also determined whether this pretreatment altered PIF4 and 5 levels the day prior to the phototropic assay (Figure 5E, F). In SUN, we observed diel regulation of PIF4 (Figure 5E) and PIF5 (Figure 5F), with a peak in the middle of the day (ZT8) and a decrease during the last hours of the day (ZT13-17). LBL had a strong effect on PIF4 protein levels (Figure 5D, E). PIF4 levels remained high for much longer during the day and only returned to the same levels as in SUN-treated samples at ZT19 (Figure 5E). LBL had a more modest effect on PIF5 protein levels which declined slightly slower in LBL than in SUN conditions (Figure 5F). As a control, we probed the membrane with CRY1 and CRY2 antibodies. As reported previously (Shalitin et al., 2002) LBL led to higher levels of CRY2 protein but not CRY1 (Figure 5E, F). We conclude that LBL has a strong effect on PIF4 protein levels particularly towards the end of the day.

**Figure 5.**
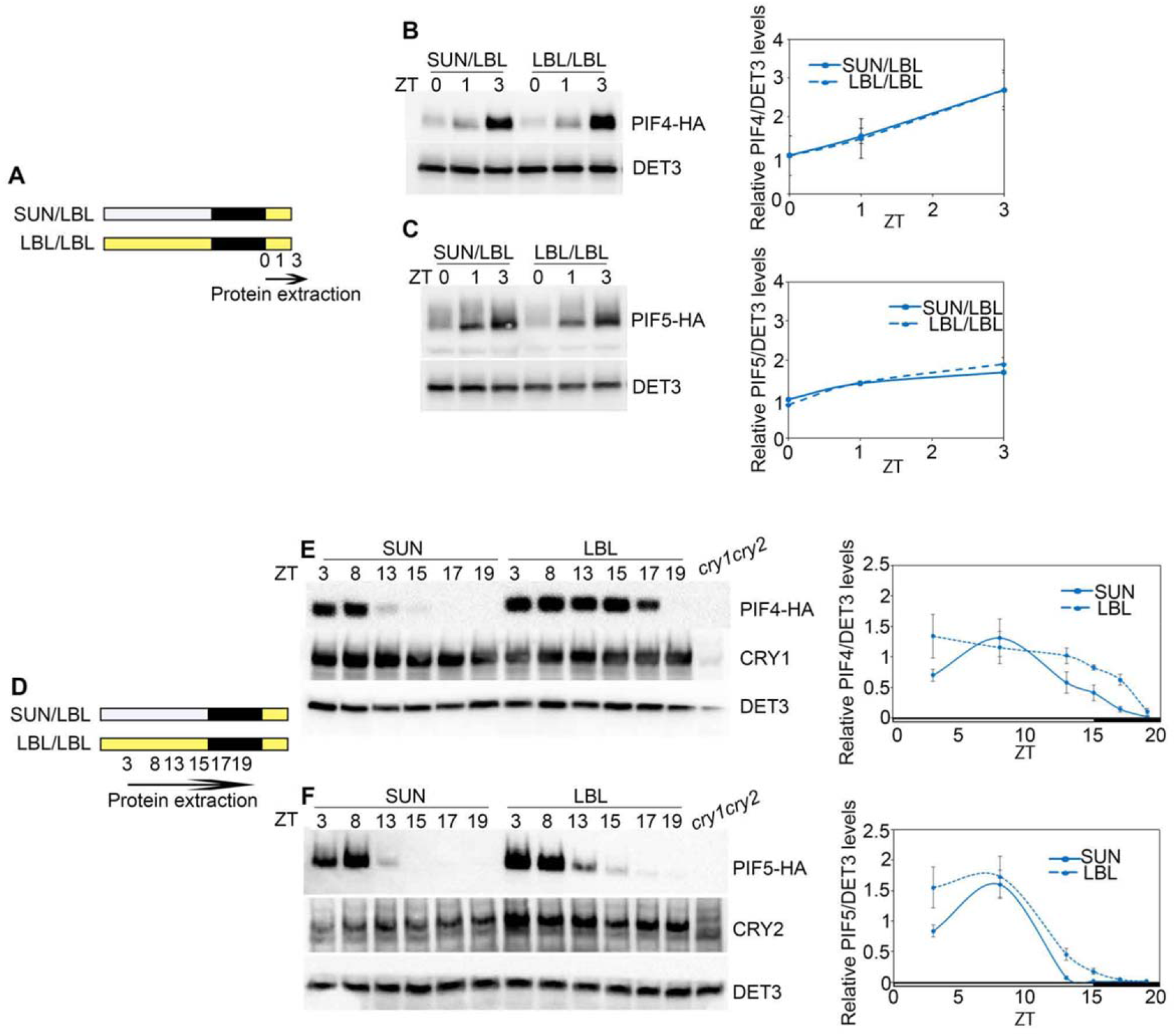
LBL leads to PIF4 and PIF5 accumulation. A and D) Schematic representations of the experiment. White boxes represent days in SUN and yellow boxes days in LBL. Black boxes represent the night. Immunoblot for PIF4-HA and PIF5-HA detected by HA antibody in *PIF4p:PIF4-HA* (B) and *PIF5p:PIF5-HA* (C) seedlings pre-treated or not with LBL at ZT0, 1, 3. Detection of PIF4-HA (E) and PIF5-HA (F) at ZT3, 8, 13, 15, 17, 19 during LBL treatment. In the immunoblot quantifications on the right, PIF4-HA and PIF5-HA levels are normalized for the loading control DET3 and they are relative to SUN ZT0 (B and C) or to SUN ZT3 (E and F) samples fixed to 1. Values are the average of three independent experiments ± SE. DET3 has used a loading control. CRY1 (E) and CRY2 (F) were detected by antibodies against the endogenous proteins.

### cry1 modulates the abundance of PIF4

To determine how LBL regulates PIF protein abundance we determined the effect of this light treatment on *PIF* transcript abundance using RT-qPCR. These experiments revealed that *PIF4* (Figure 6A), but not *PIF5* (Supplemental Figure S3), transcript levels increased in LBL, as described previously (Pedmale et al., 2016). However, the LBL treatment did not alter the diel expression profile of *PIF4* and *PIF5* (Figure 6A, Supplemental Figure S3). *PIF4* levels were higher in SUN-grown *cry1* mutants, which expressed *PIF4* levels similar to those observed in LBL-grown WT seedlings (Figure 6A). The negative effect of cry1 on PIF4 abundance was also observed by immunoblotting using a PIF4 antibody (Figure 6B). This effect on PIF4 protein abundance was confirmed and quantified comparing *PIF4p:PIF4-HA* in the WT vs. *cry* mutant background. This experiment showed that PIF4-HA levels were higher in *cry1* and *cry1cry2* particularly in SUN conditions (Supplemental Figure S4A). Moreover, PIF4-HA levels were not altered in etiolated *cry1* mutants (Supplemental Figure S4B), indicating that cry1 regulates PIF4 levels in response to blue light. The *PIF4p:PIF4-HA* line expressed higher levels of PIF4 than the WT (Supplemental Figure S4C). This provided us with an opportunity to test whether higher PIF4 levels were sufficient to promote phototropism. Interestingly, the phototropic response of *cry1*, *PIF4p:PIF4-HA* and *PIF4p:PIF4-HAcry1* was very similar, with enhanced bending compared to the WT in SUN/LBL conditions and no additive effects observed in *PIF4p:PIF4-HA cry1* (Supplemental Figure S4D). This indicates that higher PIF4 levels as observed in *cry1* or *PIF4p:PIF4-HA* promoted phototropism in SUN/LBL but further increasing PIF4 levels as in *PIF4p:PIF4-HAcry1* did not further enhance the bending response. Overexpression of PIF5 under the control of 35S promoter also enhanced phototropism in SUN/LBL conditions (Supplemental Figure S4F). Taken together these data underline the importance of PIF4 and PIF5 levels in the control of cry1-modulated phototropism.

**Figure 6.**
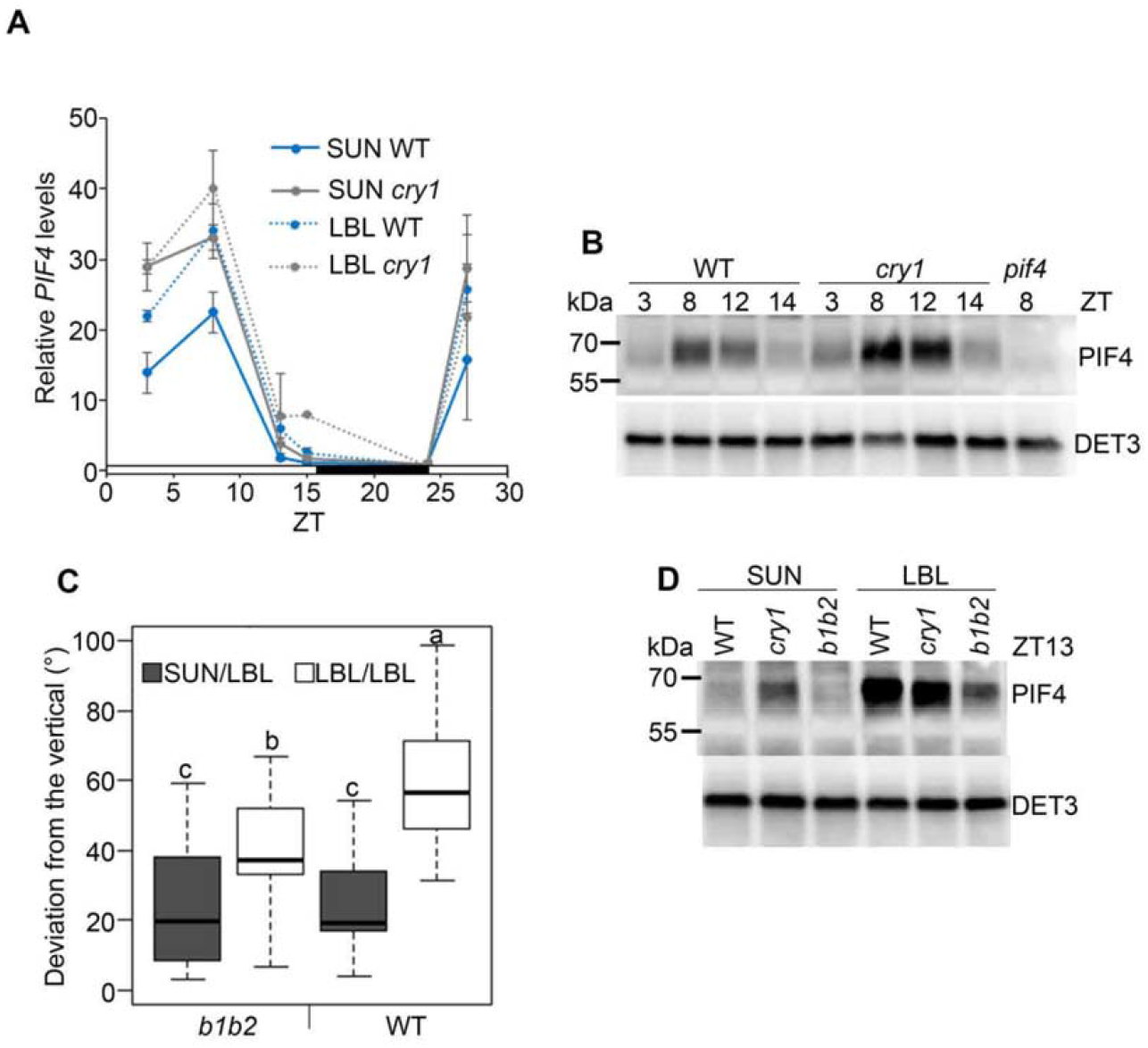
cry1 is involved in the regulation of PIF4 levels. A) RT-qPCR analysis for *PIF4* in 4-day old seedlings kept in SUN or moved to LBL at ZT0. RNA was extracted at ZT3, 8, 13, 15, 24, 27 from WT and *cry1* seedlings. Values represent the average of two independent experiments ±SE. B) Immunoblot for protein extracted from WT and *cry1* 4-day old seedlings grown in SUN at the indicated hours during the day. *pif4* mutant sample at ZT8 was used to check the specificity of PIF4 band. DET3 was used as loading control. C) Phototropic assay of WT and *bic1bic2* (*b1b2*) seedlings. Measurements were done in 3-day old 6h after lateral blue light application. Letters indicate statistically significant differences at p-value < 0.05 obtained by two way ANOVA followed by the post hoc Tukey’s HSD (n≥25). D) Immunoblot for endogenous PIF4 levels in samples collected at ZT13 kept in SUN or moved to LBL at ZT0 in. DET3 was used as loading control.

Our experiments indicate that high cry1 activity and low PIF4 levels limit phototropism in high light (SUN) conditions. A prediction of this model is that a mutant with high cry activity is expected to have reduced PIF4 levels and be less responsive to blue light gradients. We tested this using the *bic1bic2* (*b1b2*) double mutant which has higher cry activity (Wang et al., 2016). *b1b2* had the same phototropic response than the WT in SUN/LBL conditions. However, in persistent LBL which strongly promotes phototropism in the WT, *b1b2* showed a reduced phototropic response and reduced levels of PIF4 (Figure 6C, D). Taken together our data indicate that cry1 controls phototropism at least in part by controlling PIF4 levels.

### Phototropism in LBL requires auxin transport, but also biosynthesis and signaling

Asymmetrical hypocotyl growth is ensured by differential auxin distribution, which is mediated by several classes of auxin transporters including PINs (Liscum et al., 2014). Consistent with these findings, the *pin3pin4pin7* triple mutant was unable to bend in all tested conditions (Figure 7A). The analysis of an epidermis specific variant of the ratiometric auxin reporter qDII-Venus (Galvan-Ampudia et al., 2019) one hour after phototropic stimulation revealed the presence of an auxin gradient across the hypocotyl both in SUN/LBL and LBL/LBL (Figure 7B). Interestingly, in persistent LBL condition, the gradient was steeper correlating with the faster phototropic response (Figures 2, 7B). Moreover, before phototropic stimulation, the hypocotyl of seedlings pre-treated for 24 hours in LBL showed a lower qDII-Venus ratio compared to seedlings kept in SUN (Figure 7C). A low qDII-Venus ratio can be caused by higher auxin levels or higher activity of the TIR1/AFB auxin receptors, and both of these aspects may explain the faster phototropic response in persistent LBL. In response to LRFR, PIF proteins promote new auxin biosynthesis trough transcriptional activation of *YUC* genes (Hornitschek et al., 2012; Li et al., 2012). *yuc2yuc5yuc8yuc9*, as well as *taa1/sav3*, were less responsive to long LBL treatments (Figure 7D), suggesting that in persistent LBL, new auxin biosynthesis is also needed for a full phototropic response. Moreover, the reduced phototropic response in persistent LBL of *tir1* (Figure 7E) and *msg2* (Figure 7F) indicate an involvement of auxin-mediated degradation of Aux/IAA proteins. We propose that LBL-enhancement of phototropism results from a steeper auxin-signaling gradient across the hypocotyl, which may result from a coordinate action on auxin synthesis, transport and/or signaling.

**Figure 7.**
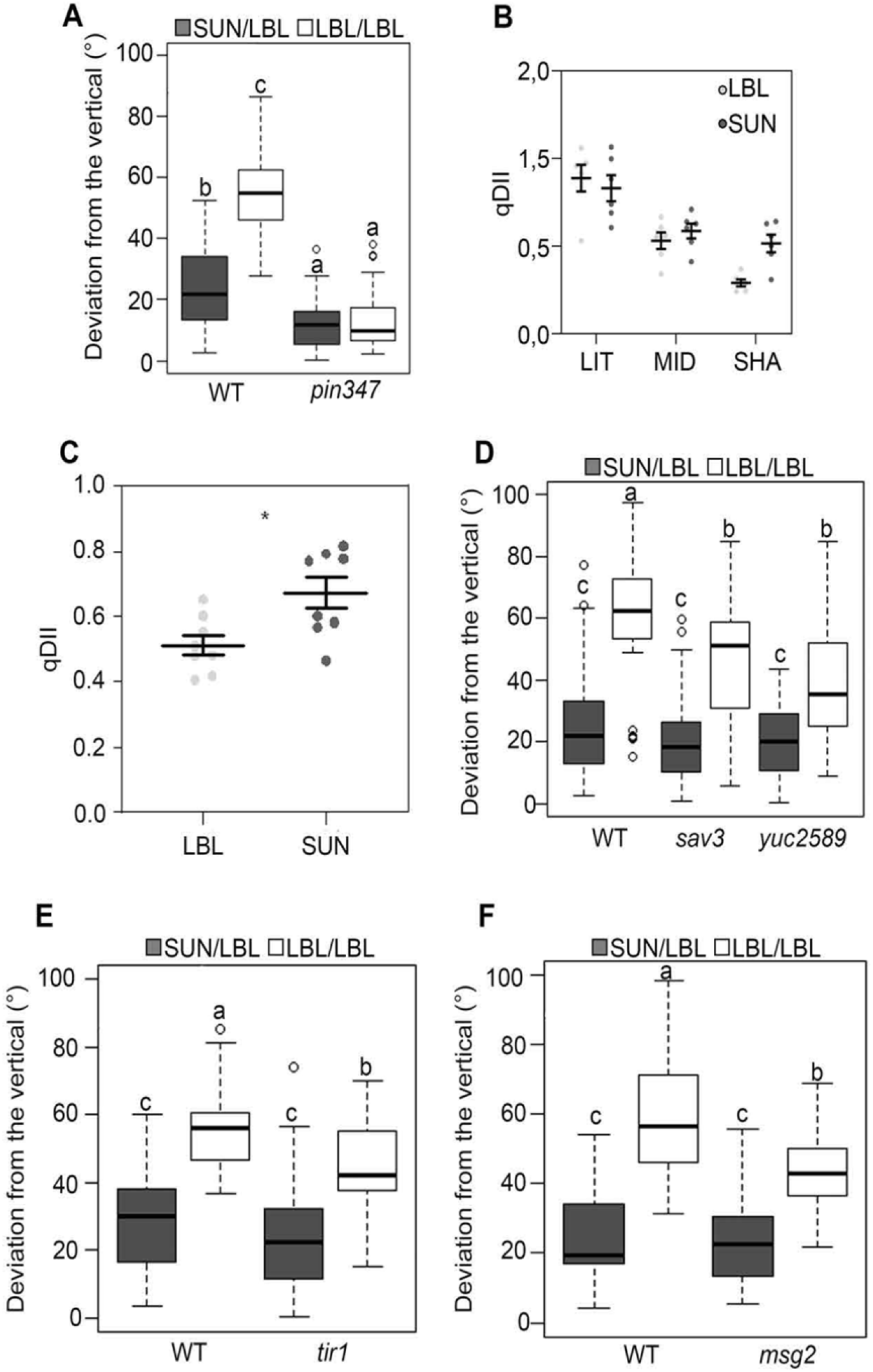
Auxin transport, biosynthesis and signaling have a role in LBL mediated phototropism. A) Phototropic assay of *pin3pin4pin7 (pin347)* mutant. B) Quantification of DII-Venus signal detected after one hour of lateral blue light application in different sides of the hypocotyl elongation zone: LIT, MID (middle), SHA (shaded). C) Quantification of DII-Venus signal detected in the epidermal cells of the hypocotyl elongation zone. Pictures were taken after 23.5-24.5 hours after the beginning of LBL treatment, or in SUN at matched ZT points. Phototropic assay of mutants for auxin biosynthesis *sav3* and *yuc2yuc5yuc8yuc9* (*yuc2589*) (D), and signaling *tir1* and *msg2* (E, F). The deviation from the vertical was measured 6h after lateral blue light application. Letters indicate statistically significant groups at p-value < 0.05 obtained by two way ANOVA followed by the post hoc Tukey’s HSD (n≥25).

## DISCUSSION

### Canopy shade promotes phototropism with a strong contribution of LBL

A positive correlation between dense vegetation and phototropism was previously documented (Ballare et al., 1992) and it was shown that the inactivation of phyB by LRFR enhances phot1-mediated hypocotyl re-orientation towards directional blue light (Goyal et al., 2016). These experiments also showed that phototropism is strongly enhanced at a very low R to FR ratio (0.2) that are typical of canopy shade and not reached prior to actual shading in neighbor proximity conditions (Ballare et al., 1990; Fiorucci and Fankhauser, 2017). We therefore investigated the effect of different features of canopy shade by testing the effects of either LRFR, LBL and the combination of both (CS) which mimics true shade in lab conditions (de Wit et al., 2016). These experiments showed that only CS leads to rapid promotion of phototropism (Figure 1B). The apparent contradiction between these results and our previous work can be explained by the very low light environment in which we performed our earlier experiments (plates were positioned in black boxes with only an opening on one side in Goyal et al., 2016). We therefore conclude that phototropism enhancement is triggered by actual vegetational shade (LRFR and LBL) rather than by neighbor proximity alone (LRFR without LBL).

Our experiments revealed that LBL strongly contributes to phototropic enhancement. For LBL to be effective on its own, it is required for several hours the day prior and during phototropic stimulation (Figure 1B, Supplemental Figure S1). This might be due to the slower effect of LBL, compared to LRFR, in promoting hypocotyl elongation (Pedmale et al., 2016). However, phototropic enhancement does not simply depend on hypocotyl elongation, given that prolonged LRFR which is highly effective in promoting hypocotyl elongation does not promote phototropism (Supplemental Figure S1). The fact that LBL alone when applied from the day prior to phototropic stimulation was sufficient to promote phototropism allowed us to specifically study the role of this component of canopy shade in phototropism enhancement. Altering blue light from the top to generate LBL also modifies the horizontal blue light gradient in our experimental setup (Figure 1A). However, several experiments allowed us to demonstrate that phototropism enhancement in LBL is not simply a consequence of a modified light gradient (Figure 1, Supplemental Figure S2). We conclude that ambient LBL is an important feature of canopy shade enhancing phototropism.

### cry1 has a negative effect on hypocotyl re-orientation of green seedlings

Our experiments show that in de-etiolated seedlings cry1 inhibits phototropism in favorable (SUN-mimicking) light conditions (Figures 2, Supplemental Figure S2). In the conditions we tested phot1 is the primary photoreceptor controlling hypocotyl reorientation and the enhanced response of *cry1* mutants depends on phot1 (Figure 3). In seedlings treated with a few hours of light to initiate de-etiolation NPH3 phosphorylation has an effect on phototropism (Sullivan et al., 2019). In our conditions, we found that NPH3 is essential for phototropism but we did not detect an effect of LBL on NPH3 phosphorylation (inferred from mobility shifts on SDS-PAGE gels) (Figure 3). Cry1-mediated phototropic suppression depends on PIF transcription factors. Under real canopy shade experiments performed outdoors the *pif4pif5pif7* triple mutant was fully epistatic over *cry1*, while the *pif4pif5* double mutant only partially suppressed *cry1* (Figure 4B). Similarly, in laboratory-simulated CS conditions we also found that *cry1* was only suppressed by the *pif4pif5pif7* triple mutant and not the *pif4pif5* double mutant (Figure 4A). However, when focusing on LBL the *pif4pif5* double mutant was sufficient to suppress *cry1* (Figure 4A), consistent with previous studies which identified PIF4 and PIF5 as the major PIFs acting downstream of cry1 in controlling shade responses (Keller et al., 2011; Pedmale et al., 2016).

### cry1 inhibits *PIF4* expression to control phototropism

The importance of PIF4 and PIF5 in controlling LBL-induced phototropism downstream of cry1 prompted us to analyze PIF4/PIF5 regulation by light and cry1. LBL treatment leads to elevated PIF4-HA and to a lesser extent PIF5-HA towards the end of the day (Figure 5). This data was confirmed for PIF4 using an antibody recognizing the endogenous protein (Figure 6). PIF4 seems to have a predominant role in regulating hypocotyl elongation in LBL, in fact, *pif4* elongates similarly to *pif4,5* double mutant, but less than *pif5* and *pif4* alone abolishes *cry1* elongation phenotype (Pedmale et al., 2016). Our data showed that cry1 regulates PIF4 levels, as shown by the analysis of PIF4 levels in *cry1* and *bic1bic2* mutants. The *cry1* mutant has higher PIF4 levels than the WT in SUN simulating conditions and the *bic1bic2* double mutant, with higher cry activity (Wang et al., 2016; Wang et al., 2017), has lower PIF4 levels in LBL correlating with a reduced phototropic response (Figure 6). The effect of LBL and *cry1* on PIF4 levels could, at least in part, be due to transcriptional regulation given that *PIF4* transcript levels are higher in LBL than in SUN conditions (Figure 6A). Moreover, LBL-regulated *PIF4* levels were essentially absent in *cry1* mutants which always expressed higher *PIF4* levels than SUN-treated WT (Figure 6A). These data are consistent with previous studies showing cry1-mediated transcriptional regulation of *PIF4* in monochromatic blue light (Ma et al., 2016; He et al., 2019). The control of PIF4 levels by cry1 is light-regulated given that we observed no effects of *cry1* on PIF4 levels in etiolated seedlings (Figure S4). Interestingly, a *PIF4p:PIF4-HA* line, which expresses higher PIF4 levels than the WT has a very similar phototropic phenotype than *cry1* without a further enhancement of the phototropic response in the *cry1 PIF4p:PIF4-HA* line (Figure S4). This suggests that high levels of PIF4 alone are sufficient to promote phototropism and that a major level of cry1 regulation is the transcriptional control of PIF4 accumulation. These observations are in agreement with previous data showing that when expressed from a constitutive promoter PIF4 protein levels are unchanged in the *cry1* mutant (Ma et al., 2016). The precise mechanism underlying cry1-mediated enhancement of *PIF4* expression remains unknown. However, it is noteworthy that cry2 modulates gene expression in a blue light-regulated fashion when expressed in a heterologous system (Yang et al., 2018). Our data do not rule out additional levels of PIF4 regulation by cry1 such as post-transcriptional regulation or inhibition of PIF4 activity (Ma et al., 2016; Pedmale et al., 2016). Yet, the striking correlation between cry1-mediated PIF4 accumulation and LBL-modulated phototropism highlights the importance of cry1-regulated PIF4 abundance at the transcriptional level.

### The importance of auxin for LBL-mediated phototropic enhancement

Several reports have demonstrated impaired hypocotyl elongation responses to LBL in mutants defective in auxin transport and auxin biosynthesis (Pierik et al., 2009; Keuskamp et al., 2011; de Wit et al., 2016). Deficient enhancement of the phototropic response by LBL in the *sav3*, *yuc2yuc5yuc8yuc9*, *pin3pin4pin7, tir1* and *msg2* mutants (Figure 7) indicate that this process requires normal auxin synthesis, transport, perception and signaling. A priori, the phenotype of these mutants might simply indicate that normal auxin synthesis, transport, perception and signaling are a condition for the LBL effects, or that the auxin system carries LBL information. In this regard, PIFs regulate auxin signaling at multiple levels including biosynthesis, transport, perception and signaling (Oh et al., 2014; Kohnen et al., 2016; Iglesias et al., 2018; Pucciariello et al., 2018) and therefore, LBL-mediated phototropic enhancement may affect more than one of these levels of regulation. For instance, shade cues (LBL and/or LRFR) promote the expression of several *PINs* including *PIN3* and *PIN7* (Keuskamp et al., 2011; Kohnen et al., 2016). Moreover, cry1 and the PIFs regulate *PIN* expression in an antagonistic way with higher expression in *cry1* mutants and reduced expression in *pif* mutants (Hornitschek et al., 2012; Li et al., 2012; He et al., 2019). Given that PIF4 and PIF5 directly bind to the promoter of *PIN3* (Hornitschek et al., 2012), it is possible that the cry1-mediated regulation of PIF4 abundance modulates the phototropic response via PIF-controlled *PIN* expression. At least under LRFR, PIFs also promote the expression of several *YUC* genes to enhance phototropism (Goyal et al., 2016). Therefore, we used qDII-Venus to investigate whether the auxin system actually carries the LBL information. Our data indicate that the latter is actually the case because a LBL pretreatment leads to higher auxin levels and/or sensitivity in the hypocotyl (Figure 7C) and a steeper gradient of auxin levels and/or sensitivity upon phototropic stimulation (Figure 7B), which correlates with enhanced phototropism (Figure 1).

We conclude that phototropic enhancement by canopy shade involves changes in activity of at least three photoreceptors: phot1, cry1 and phyB (Figures 2, 3) (Goyal et al., 2016). In shade, the reduced activity of cry1 and phyB permits enhanced PIF abundance, leading to a modification of auxin signaling status in the hypocotyl to promote phototropism when phot1 perceives the blue-light gradient (Figure 7) (Goyal et al., 2016).

## MATERIALS AND METHODS

### Plant material

The following *Arabidopsis thaliana* (Col-0 ecotype) mutants were previously characterized: *cry1-304* (Mockler et al., 1999)*; phot1-5* (Huala et al., 1997)*; nph3-6* (Motchoulski and Liscum, 1999)*; cry2-1* (Guo et al., 1998)*; cry1-304cry2-1; cry1-304pif4-101pif5-3, cry1-340pif4-301pif5-3pif7-1* and *cry1phyB* (Fiorucci et al., 2019)*, phyB-9; phyB-9pif4-101pif5-3* and *phyB-9pif4-101pif5-3pif7-1* (Goyal et al., 2016)*, pif4-101pif5-3 and OXPIF5 (Lorrain et al., 2008)*; *pif4-101pif5-3pif7-1* (de Wit et al., 2015)*; PIF4p:PIF4-HApif4-301* (Galvao et al., 2019) and *PIF5p:PIF5-HApif5-3* (de Wit et al., 2016)*, bic1bic2* (Wang et al., 2016)*, sav3/taa1* (Tao et al., 2008)*, yuc2yuc5yuc8yuc9* (Kohnen et al., 2016)*, tir1-1*(Ruegger et al., 1998)*, msg2* (Tatematsu et al., 2004)*, pin347* (Willige et al., 2013). *pPIF4:PIF4-HApif4-101cry1-304, pPIF4:PIF4-HApif4-101cry1-304cry2-1, cry1-304phot1-5* were obtained by crosses and the primers used for genotyping are listed in Supplemental Table1. The epidermis specific promoter PDF1 (protodermal factor 1) (Abe et al., 2001) driving the expression of a DII-VENUS-N7-2A-TagBFP-sv40 (qDII) (Galvan-Ampudia et al., 2019) was assembled by Gateway technology (Invitrogen) and transformed in *Arabidopsis thaliana* (Col-0).

### Growth and light conditions

Seeds were surface sterilized according to Kohnen et al., 2016 and placed on plates containing half-strength Murashige and Skoog medium (½ MS), 0.8% (w/v) phytoagar (Agar-Agar, plant; Roth) and MES. After 2 days of stratification (4°C and dark), seedlings grew inside customized black boxes. In order to avoid varaibility due to the light gradient towards the bottom of the black boxes during seedlings growth, we sowed 32 seeds only in the upper part of the plate arranged in 4 rows. Seedlings were grown in long day (16h day/8h night, 21°C/19°C) in presence of 95μmol m^−2^s^−1^ of PAR (Photosynthetically Active Radiation) measured by white diffuser filter combined with PAR filter (Radiometer Model IL1400A, https://www.intl-lighttech.com/product-group/light-measurement-optical-filters). The single layer of yellow filter (010 Medium Yellow, LEE Filters) used to cover up the seedlings cut down blue light from 7μmol m^−2^s^−1^ (SUN) to 0.7μmol m^−2^s^−1^ (LBL) and it was measured by white diffuser filter combined with blue filter (400-500nm). The R (640–700 nm)/FR (700–760 nm) ratio was measured using the Ocean Optics USB2000+ spectrometer. In SUN and LBL the R/FR ratio was 1.25 and in LRFR and CS it was 0.3, obtained by adding FR LED to white light lamps. The LEE filter number 298 0.15ND was used in Figure Supplemental S1 to reduce PAR similarly to the yellow filter used for LBL treatment. Transmission through ND filter is 69.3%, the transmission of PAR through the yellow filter is about 76.6%.

### Phototropism

Phototropic stimulation by the application of lateral blue light was always started after the lights turned on, between ZT0 and ZT0.5 by removing one side of the black boxes and supplying unilateral blue light by LED. The LED source was placed 60 cm distant from the black boxes to raise the horizontal blue light up to 8μmol m^−2^s^−1^. In addition, we calculated the blue light differential between the illuminated side and the top (i.e., the light coming from above), because the yellow filter in the LBL condition affected this differential and we had to evaluate its potential physiological impact. The log10 of the side-to-top differential of blue light was 0.0 (log10 (side blue light (8μmol m^−2^s^−1^) - top blue light (7μmol m^−2^s^−1^))= 0) under SUN (high blue light from above) and 0.9 (log10 (side blue light (8μmol m^−2^s^−1^) - top blue light (0,7μmol m^−2^s^−1^))= 0.9) under LBL. Only in the SUN condition presented in Figure Supplemental S2B, we increased the side blue light up to 24μmol m^−2^s^−1^ to obtain a log10 of the differential between the illuminated side and the top similar to that observed under LBL conditions (log10 (side blue light (24μmol m^−2^s^−1^) −top blue light (12μmol m^−2^s^−1^)=1.0).

### Measurement of the phototropic response

Pictures taken before and after phototropism were analyzed with a MATLAB script developed in Fankhauser’s lab to obtain bending angle and hypocotyl length values. The bending angle was calculated as the deviation from the vertical of the upper part of the hypocotyl (75-95% of the hypocotyl length). Since both *phot1cry1* and *phot1* are impaired in their gravitropic response, we straightened the seedlings before phototropism and we calculated the bending angle between T0 and T6 hours of treatment. For the creation of box plots and to computer the two-way ANOVA (aov) and computed Tukey’s Honest Significance differences (HSD.test), the [agricolae package] of the R software was used. Similar results were obtained in three independent experiments.

### RNA extraction and gene expression analysis

20-25 seedlings grown in horizontal Petri dishes were frozen in liquid nitrogen and total RNA was extracted with the RNeasy Mini Kit Qiagen®. 500ngr of RNA were used for cDNA preparation using Superscript™ Transcriptase (Invitrogen). The qPCR was performed as described in Kohnenet al., 2016. Relative gene expression was calculated as following: (Eff_(TARGET)_)^ΔCtTARGET^/(Eff_(HK)_)^ΔCtHK^. In the calculation the average of the Ct values derived from two housekeeping genes (*UBC* and *YSL8*) was used to normalize the expression of the target gene. The primers sequences are listed in Supplemental Table1.

### Protein extraction and immunoblot analysis

Protein extraction for the detection of HA (1:2000), CRY1 (1:4000 anti-rabbit secondary antibody), CRY2 (1:3000 anti-rabbit secondary antibody) and NPH3 (1:3000 anti-rabbit secondary antibody (Motchoulski and Liscum, 1999)) was performed according to Galvao et al., 2019. Protein extraction for PIF4 N3 (1:3000, Abiocode R2534-4) detection was performed according to Fiorucci et al., 2019. Samples extracted from 20-25 seedlings were separated on 4-20% MiniProtean TGX gels (BIO-RAD), except for the NPH3 immunoblot, for which 8% Acrylamide SDS-PAGE gels were used. All samples were transferred to Nitrocellulose membrane with the Trans-Blot Turbo RTA transfer kit (BIO-RAD Cat.#170-4270). The chemiluminescent signal was detected with Immobilon Western Chemiluminescent HRP Substrate (Millipore) on an ImageQuant LAS 4000 mini (GE Healthcare). The HA/DET3 signals were quantified using the ImageJ software.

### DII-Venus detection and quantification

The genotype used was pPDF1∷DII-n7-Venus-2A-mTurquoise-sv40 t35 in the Col-0 background. Seedlings were grown as for phototropic experiment. On the fourth day, half of the plates were shifted in LBL and half of them were kept in SUN. The fifth day, all plates were covered with a yellow filter before the start of the day and 1h after the start of the day one side of the black box was opened to perform the phototropism assay. Confocal images were taken between ZT23,5 and ZT0,5 and between 1h and 2h after starting the phototropic assay. All pictures were taken in the epidermis in the elongation zone using an LSM710 confocal microscope (Zeiss) equipped with an EC Plan-Neofluar 20x/0.50 objective. For Venus we used an Argon laser (514nm) and detected between 519 and 559nm. For BFP we used a 405nm diode laser and detected between 430 and 470nm. Image analysis was performed in ImageJ. A threshold was applied to the BFP channel to segment the nuclei. Mean intensity was measured inside each nucleus in the BFP and the Venus channels and qDII was calculated as the ratio between these two values for each nucleus. Each data point represents the average of 3-4 seedlings coming from one plate, and at least 5 nuclei were measured in each seedling.

## ACKNOWLEDGEMENTS

We would like to thank Anne-Sophie Fiorucci for providing the R script for boxplot representations. Martine Trevisan for technical assistance. The GTF facility at University of Lausanne for qPCR assistance. The CIF facility at the University of Lausanne for assistance with confocal microscopy. Chentao Lin for CRY1 and CRY2 antibodies. Géraldine Brunoud for her help with the design of the qDII-VENUS sensor. All the Fankhauser lab for the fruitful discussions.

## Supplemental Material

- **Supplemental Figures S1 to S4**
- **Supplemental Table 1**. List of Primers

